# Aberrant striatal oscillations after dopamine loss in parkinsonian non-human primates

**DOI:** 10.1101/650770

**Authors:** Arun Singh, Stella M. Papa

## Abstract

Dopamine depletion in Parkinson’s disease (PD) is associated with abnormal oscillatory activity in the cortico-basal ganglia network. However, the oscillatory pattern of striatal neurons in PD remains poorly defined. Here, we analyzed the local field potentials in one untreated and five MPTP-treated non-human primates (NHP) to model advanced PD. Augmented oscillatory activity in the alpha (8-13 Hz) and low-beta (13-20 Hz) frequency bands was found in the striatum in parallel to the motor cortex and globus pallidus of the NHP-PD model. The coherence analysis showed increased connectivity in the cortico-striatal and striato-pallidal pathways at alpha and low-beta frequency bands, confirming the presence of abnormal 8-20 Hz activity in the cortico-basal ganglia network. The acute L-Dopa injection that induced a clear motor response normalized the amplified 8-20 Hz oscillations. These findings indicate that pathological striatal oscillations at alpha and low-beta bands are concordant with the basal ganglia network changes after dopamine depletion, and thereby support a key role of the striatum in the generation of parkinsonian motor abnormalities.

## Introduction

Abnormal oscillatory activities in the basal ganglia network are correlated with motor abnormality in Parkinson’s disease (PD) (Brown, 2003; Singh, 2018). The degeneration of nigral dopaminergic neurons that project to the striatum is the pathological hallmark of PD (Damier *et al.*, 1999), but the role of the striatum as an important source of abnormal activity patterns (spiking and oscillations) in PD has remained unclear. Our previous studies have shown aberrant striatal neuronal spiking activity in parkinsonian non-human primates (NHPs) and patients with PD (Liang *et al.*, 2008; Singh *et al.*, 2015; Singh *et al.*, 2016). Indeed, chronic dopamine loss leads to major changes in the firing of striatal projection neurons, i.e. hyperactivity of >10-fold increase in mean firing rate compared to normal state, and unstable responses to dopamine. These significant changes in the SPN spiking are likely to be accompanied by altered local inputs and oscillatory activity. In extrastriatal basal ganglia structures, a consistent increased oscillatory activity has been found in animal models and patients with PD (Hutchison *et al.*, 2004; Devergnas *et al.*, 2014; de Hemptinne *et al.*, 2015). A series of studies led to establish the increased 8-30 Hz oscillations in the cortico-basal ganglia network as a key feature of the pathophysiology of PD (Stein and Bar-Gad, 2013). Treatment of PD by high frequency deep brain stimulation or L-Dopa that improves motor disability also suppresses the abnormally synchronized 8-35 Hz oscillations in the basal ganglia (Brown *et al.*, 2001; Kuhn *et al.*, 2006).

In contrast to the more typical 13-30 Hz (beta-band) found in human studies, NHP recordings detected synchronized basal ganglia oscillations in the 8-13 Hz frequency band, and suggested that these rhythms may be a subclass of, or similar to, the beta band observed in PD patients (Stein and Bar-Gad, 2013; Devergnas *et al.*, 2014). The improvement of parkinsonian symptoms also correlates with a decrease in 8-13 Hz oscillations in the cortico-basal ganglia network of Parkinson’s disease (Brown, 2007). However, the function of oscillations in this lower frequency band (8-13 Hz) in the mechanisms of motor dysfunction in PD remains unknown. Furthermore, the source of these oscillations in the basal ganglia network has not been revealed in the previous studies. Thus, the oscillatory pattern of the striatum, the basal ganglia recipient structure, after dopamine denervation becomes an essential piece of information to understand the role of various pathological network oscillations in PD.

Here, we examined striatal and other basal ganglia nuclei for oscillatory activities in normal and parkinsonian NHPs. In addition, we analyzed the striatal oscillations before and after acute L-Dopa administration in correlation with improvement of motor deficits. This study shows that abnormal oscillations in the alpha (8-13 Hz) and low-beta (13-20 Hz) bands are present in the striatum of parkinsonian primates, and that these striatal oscillations are modulated by dopamine replacement.

## Material and Methods

### Animals and MPTP Treatment

Six adult macaques (four female rhesus monkeys, *Macaca Mulatta*: Gl, Va, Na, Sa; and two male cynomolgus monkeys, *Macaca Fascicularis*: Br and Ch; all 5-9 kg) were used in this study. All experimental protocols were performed in accordance with the *National Institutes of Health Guide for the Care and Use of Laboratory Animals* (1996). Five animals received weekly intravenous injections of MPTP (0.2-0.6 mg/kg) until motor disability stabilized at moderate parkinsonism, as measured by the standardized Motor Disability scale for MPTP-treated primates (Potts *et al.*, 2014; Singh *et al.*, 2015). One animal (Sa) was not treated and used as a normal control. Parkinsonian rhesus NHPs were previously treated with daily L-Dopa regimen (Sinemet®, 100-200 mg/day), but parkinsonian cynomolgus NHPs had not been exposed to L-Dopa. For acute L-Dopa subcutaneous injections (L-Dopa methyl ester plus benserazide) during the recordings of parkinsonian cynomolgus NHPs (not chronically exposed to L-Dopa), we selected doses based on our previous recordings of moderately parkinsonian primates to induce a response compatible with the “on” state (reversal of parkinsonism) and the restraint of the animal in the primate chair.

### Surgical Procedures and LFP Recordings

Stainless steel recording chambers were stereotaxically implanted in the coronal plane with a 15-20° angle in all NHPs to allow the trajectory of recording electrodes to include the motor cortex, the striatum and the internal pallidum (GPi). Implant surgeries were conducted under general anesthesia in all animals. The electrophysiological mapping of the caudate, putamen, and GPi was performed by extracellular recordings with tungsten microelectrodes (FHC, Bowdoinham, ME; impedance = 0.1–0.3 MΩ at 1 kHz) using standard methods (DeLong, 1971; Liang *et al.*, 2008). Subsequently, the regional LFPs were recorded using the same tungsten microelectrodes and the recording chamber implanted on the skull or the connection to the amplified electrical ground as reference. To validate the reliability of the striatal LFP signals, we performed additional recordings with a reference electrode in the striatum for offline reconstruction of bipolar LFPs for comparison with unipolar striatal LFP. In bipolar recordings, the LFP signal was similar to that recorded with the standard unipolar method (see Fig. 1A and C).

**Figure 1.**
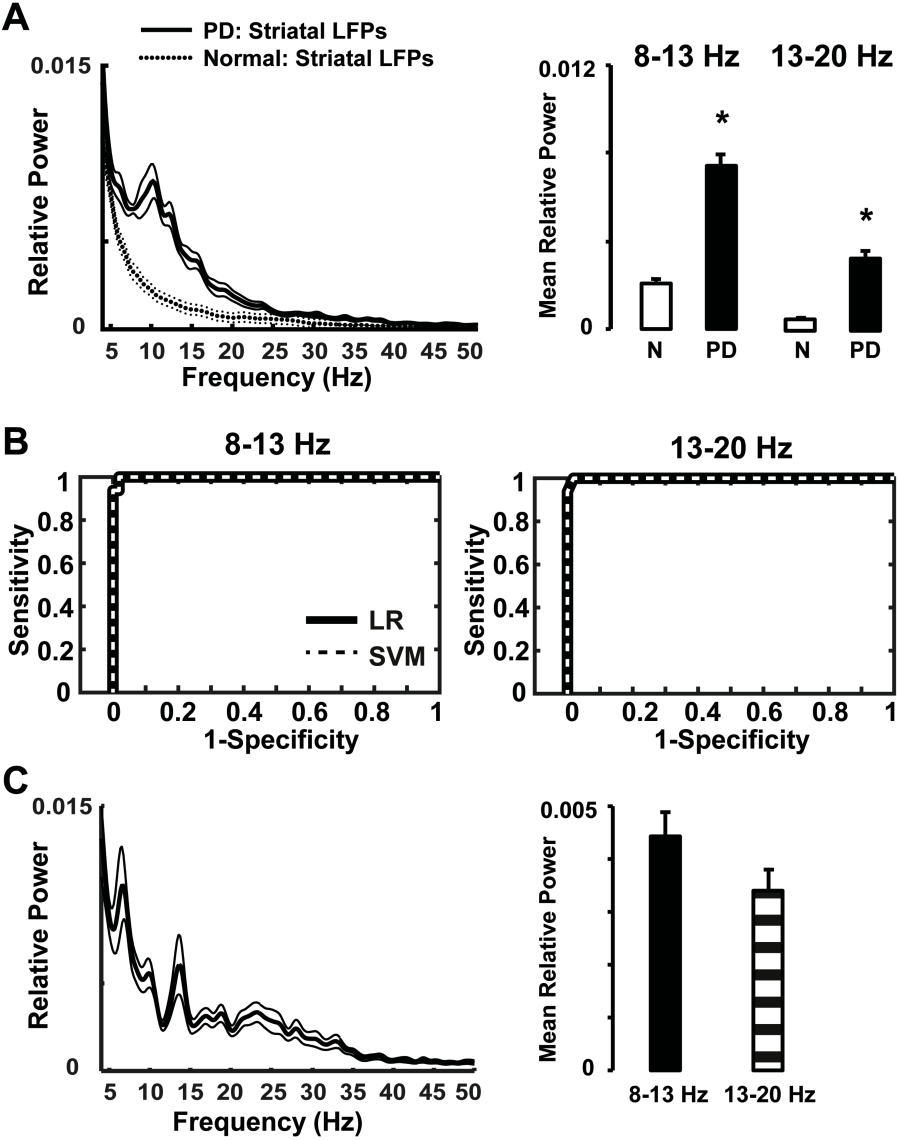
Striatal local field potentials (LFPs) in parkinsonian primates. (A) Mean relative power in the 8-13 Hz and 13-20 Hz frequency bands was significantly increased in the dopamine depleted striatum of NHPs as compared with the intact striatum in the normal NHP. (B) Classification of control versus parkinsonian NHPs based on power values of 8-13 Hz and 13-20 Hz frequency bands. Receiver operating characteristic plots show the true (sensitivity) vs. false positive rates (1-specificity) of normal vs. MPTP-treated NHPs discrimination for each frequency band separately using logistic regression (LR) and support vector machines (SVM) methods. (C) Bipolar striatal LFPs in a MPTP-treated NHP. Spectral plot and relative power in the 8-13 Hz and 13-20 Hz frequency bands confirm the presence of the same oscillatory activities as shown by monopolar LFP signals. Thick and thin lines represent mean and standard deviation, respectively. Significant difference is denoted by \**P*<0.01.

LFPs were recorded in one normal NHP (Sa) and in three parkinsonian NHPs (Va, Na, and Gl) during the ‘off’ state (basal parkinsonian motor disability) in the motor cortex, striatum, and GPi. To determine the presence of coherent oscillatory activity within the cortico-basal ganglia network, LFP signals were simultaneously collected in two network stations, i.e. (i) cortical and striatal regions, and (ii) striatal and GPi regions in one parkinsonian NHP (Va). Striatal LFPs were recorded in two parkinsonian NHPs (Br and Ch) during the “off” and “on” states from putamen. Typically, the ‘on’ state was evidenced by behavioral changes approximately 20 minutes after the acute systemic injection of L-Dopa. LFP signals (sampling frequency: 2000 Hz, Blackrock Microsystems; sampling frequency: 20 kHz, Multichannel Acquisition Processor, Plexon Inc.) were amplified and low pass filtered with 750 Hz. LFPs were recorded continuously for at least 3-4 min in each motor states. Data were monitored online to ensure correlation with motor states. Collected LFPs were stored for offline analysis.

### Data Analysis

Continuous LFP data were analyzed using Matlab software (Math Works, Natick, MA), with custom codes on the basis of standardized signal analysis functions (signal analysis toolbox). Power spectral analysis of LFP signals was computed applying the “pwelch” method. All data were resampled to 1024 Hz and band passed 1-90 Hz (Butterworth zero phase filter). A 60 Hz notch filter was also used to eliminate line-noise. Visual examination was initially performed to remove time periods with obvious artifacts. Signals were segmented into 5 sec epochs, which were transformed into the power spectrum domain (pwelch method: 1024 samples window size overlapped by 50%). A frequency range of 1-90 Hz was selected to compute relative power spectrum to reduce the inter-recording variation. Subsequently, all spectra were averaged. Previous studies of basal ganglia oscillation in MPTP-treated NHPs detected increases in the 8-20 Hz frequency range (Stein and Bar-Gad, 2013; Devergnas *et al.*, 2014). As we observed two distinct spectral peaks in the 8-13 Hz and 13-20 Hz frequency ranges, we computed the mean relative LFP-power at these two frequency bands. We used 4-50 Hz range for visualizing the relative power spectrum. Coherence analysis between cortico-striatal and striato-pallidal LFPs was computed using the “csd” matlab function with a window size of 1024, which provided a frequency resolution of ∼1 Hz with 50% overlap. Differences between relative power of striatal LFPs (8-13 Hz and 13-20 Hz) recorded in normal and parkinsonian NHPs were analyzed with unpaired t-test. Differences between “off” and “on” states for 8-13 Hz and 13-20 Hz frequency bands were analyzed with paired t-tests.

The relative power of 8-13 Hz and 13-20 Hz frequency band oscillations were computed to differentiate normal from parkinsonian NHPs, and “off” from “on” motor states using logistic regression (LR). We examined the accuracy of the relation (predictor or classifier) using a two-class problem in the support vector machine (SVM) method on the average of 10 cross validation tests on 8-13 Hz and 13-20 Hz frequency bands values. Data were divided into ten groups and then trained with nine groups and tested with one group. This was repeated ten times, while each group was used as a test group. Afterwards, single performance estimation was calculated. We used ‘svmtrain’ and ‘svmclassify’ matlab functions for classification and ‘classperf’ matlab function to estimate the performance or accuracy of the classifier. Accuracy of the classifier was displayed using a Receiver Operating Characteristic (ROC) curve, a graphical plot of the sensitivity vs. (1-specificity).

## Results

In the normal NHP, there is no predominant striatal oscillatory activity in any frequency range. In contrast, dopamine loss in MPTP-treated NHPs induces aberrant striatal oscillations with relative power at 8-13 Hz and 13-20 Hz frequency bands as compared to the normal condition (*P* = 0.001; Fig. 1A). Further analysis of the relation between the relative power of 8-13 Hz and 13-20 Hz frequency bands to normal and parkinsonian conditions in NHPs revealed a strong correlation with notable sensitivity (8-13 Hz: 99 % LR method, 99 % SVM method; 13-20 Hz: 99 % LR method, 99 % SVM method; Fig. 1B), and accuracy (8-13 Hz: 97 %; 13-20 Hz: 98 %) for these parameters to classify control versus MPTP-treated NHPs. These results were highly constant across algorithms and cross-validation procedures.

In parkinsonian NHPs, the relative power of 8-13 Hz and 13-20 Hz frequency bands in the striatum was similar to those recorded in the motor cortex and GPi (Fig. 2). In the normal NHP, fixed electrodes in the striatum prevented recordings of LFPs in the cortex and GPi. LFPs recorded simultaneously in the motor cortex and the striatum or the striatum and GPi show increased coherence in the 8-13 Hz and 13-20 Hz frequency bands in parkinsonian NHPs (Fig. 3). Similarly, previous human and rodent studies have demonstrated an increase in coherence at same frequency bands between cortical and basal ganglia regions in the parkinsonian state (Sharott *et al.*, 2005; Litvak *et al.*, 2011; Shimamoto *et al.*, 2013). Overall, these results confirm that synchronized striatal oscillations in the 8-20 Hz frequency band can reflect a network pattern that occurs after dopamine loss.

**Figure 2.**
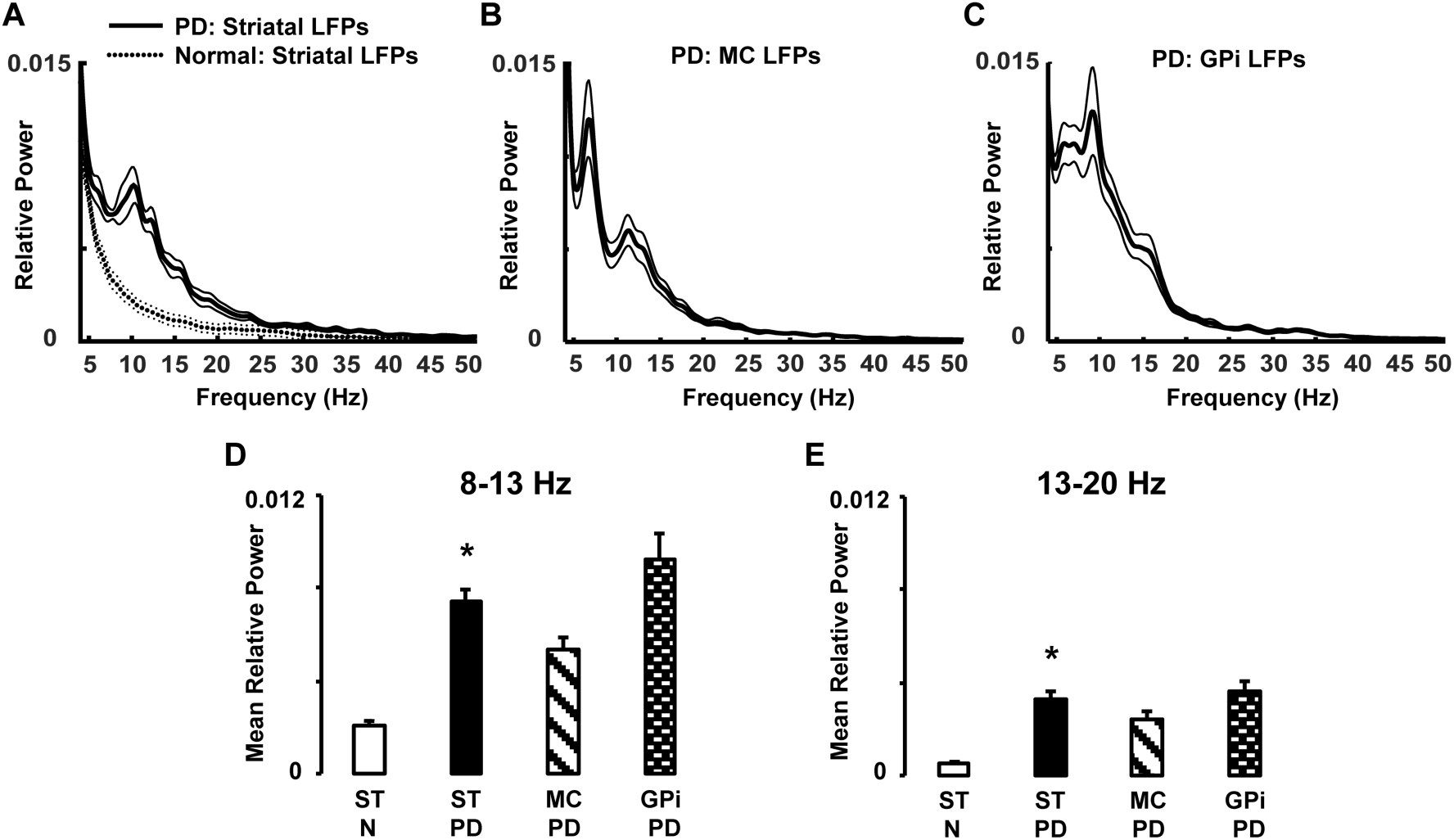
Local field potentials (LFPs) recorded from motor cortex and basal ganglia nuclei. (A). Striatal LFPs collected in normal and parkinsonian NHPs. (B and C) Cortical and pallidal (GPi) oscillations recorded in parkinsonian NHPs. (D-E) Comparison of the relative power of 8-13 Hz and 13-20 Hz frequency bands in each of the studied regions. In the parkinsonian state, the relative power of LFPs at 8-13 Hz and 13-20 Hz was similar across the motor cortex, the striatum and the GPi. Thick and thin lines represent mean and standard deviation, respectively. Significant difference is denoted by \**P*<0.01. N= Normal NHP; PD = MPTP-treated NHP; MC= motor cortex; ST= striatum; GPi= globus pallidus internus.

**Figure 3.**
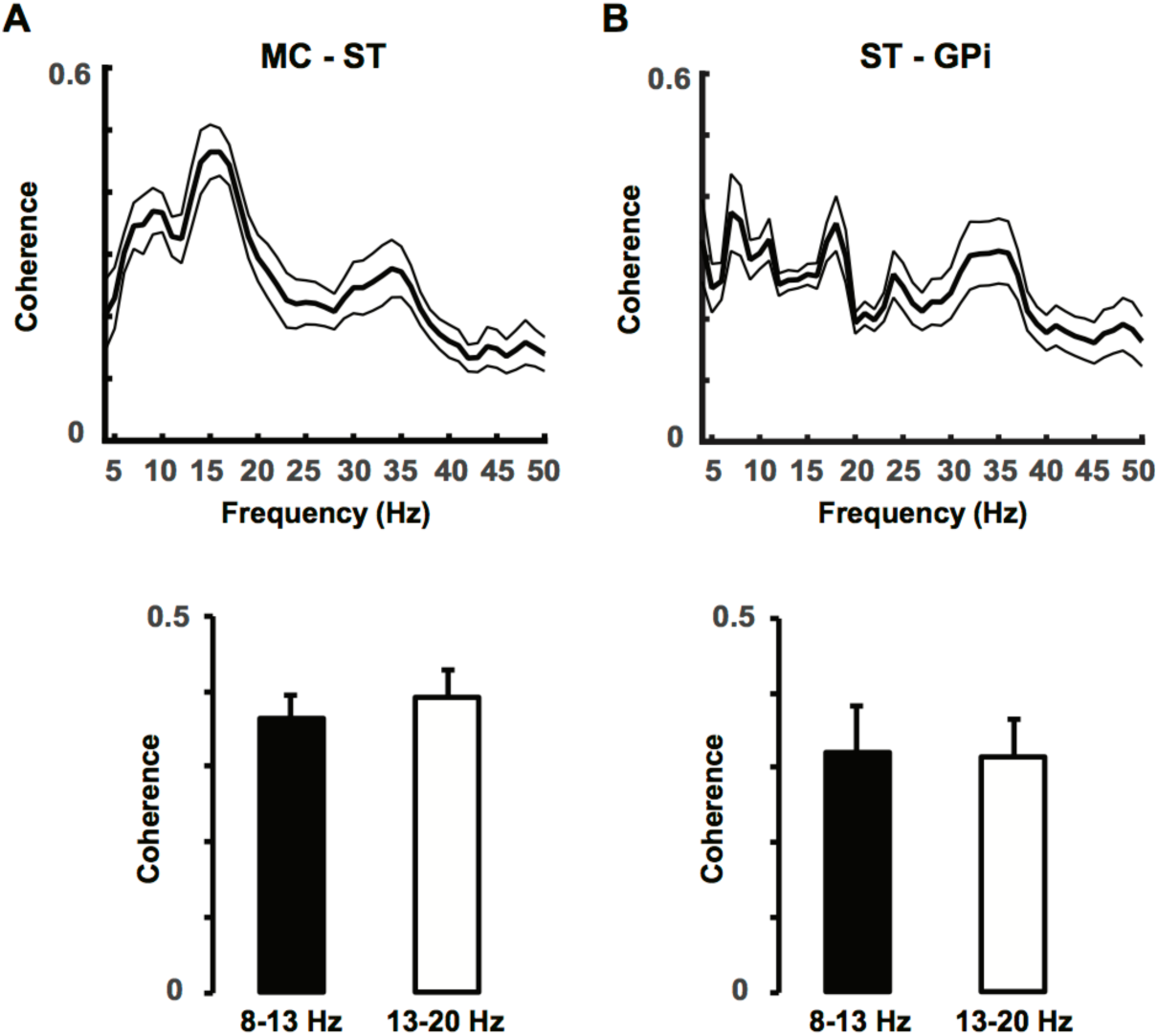
Coherence of oscillatory activity in the cortico-striato-pallidal network in parkinsonian NHPs. (A) Increased coherence of LFPs between the motor cortex and the striatum similarly shown at both 8-13 Hz and 13-20 Hz frequency bands (bottom graph). (B) Increased coherence of LFPs also between the striatum and the GPi at both frequency bands. These results demonstrate the presence of augmented oscillatory activity at 8-20 Hz in the basal ganglia circuits after dopamine loss. Thick and thin lines represent mean and standard deviation, respectively. MC=motor cortex; ST=striatum; GPi=globus pallidus internus.

Power spectral analysis of the striatal LFPs recorded in NHPs after acute L-Dopa injection inducing a motor response (“on” state) revealed markedly reduced relative power in the 8-13 Hz (*P* = 0.001) and 13-20 Hz (*P* = 0.01) frequency bands (Fig. 4A and B). Both changes in 8-13 Hz and 13-20 Hz frequency bands following dopamine replacement were highly sensitive (8-13 Hz: 71 % LR method, 71 % SVM method; 13-20 Hz: 68 % LR method, 68 % SVM method; Fig. 4C) and accurate (8-13 Hz: 70 %; 13-20 Hz: 66 %) to distinguish “off” and “on” states of MPTP-treated NHPs. These results are in line with the previous human and animal studies that have shown reduced alpha and beta-band power in extrastriatal regions after L-Dopa administration (Brown *et al.*, 2001; Kuhn *et al.*, 2006; Devergnas *et al.*, 2014).

**Figure 4.**
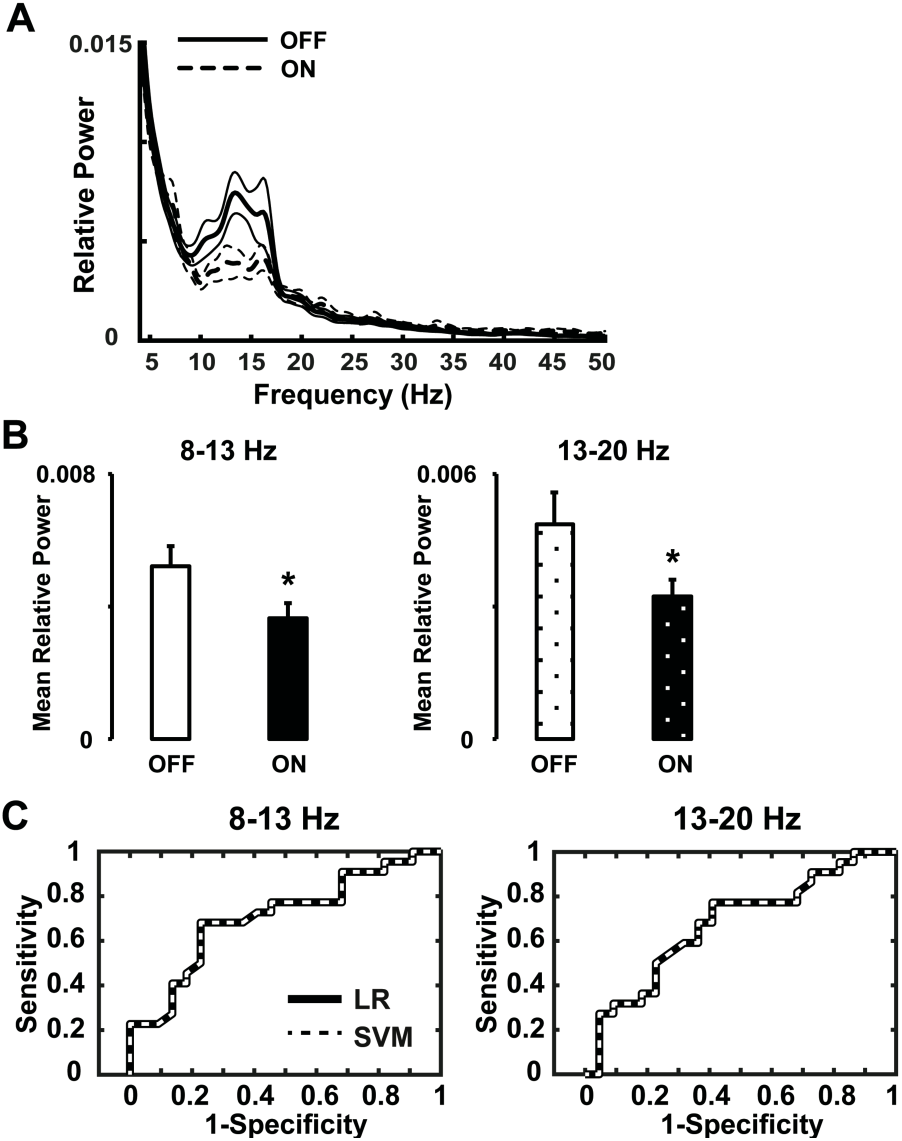
Striatal LFPs before and after acute L-Dopa administration in parkinsonian NHPs. (A and B) Relative power of the striatal oscillations at 8-13 Hz and 13-20 Hz frequency range was significantly decreased after L-Dopa injection. (C) Classification of “off” versus “on” states in parkinsonian NHPs based on power values of 8-13 Hz and 13-20 Hz frequency bands. Receiver operating characteristic plots show the true (sensitivity) vs. false positive rates (1-specificity) of “off” vs. “on” state discrimination for each frequency band separately using logistic regression (LR) and support vector machines (SVM) methods. Thick and thin lines represent mean and standard deviation, respectively. Significant difference is denoted by \**P*<0.01.

## Discussion

Current results showed the presence of augmented 8-13 Hz and 13-20 striatal oscillations in MPTP-treated NHPs after dopamine depletion. Similarly, acute reduction of dopamine in the striatum amplifies striatal beta oscillations in rodents (Costa *et al.*, 2006). These data also extend the findings of previous studies to demonstrate the changes in extrastriatal basal ganglia oscillatory activity in the 8-13 Hz and 13-20 Hz frequency bands in levodopa-treated parkinsonian animals (Stein and Bar-Gad, 2013; Devergnas *et al.*, 2014). Different cell types in the striatum can be the main cause of induction of abnormal filed potentials after dopamine loss. Robust 8-30 Hz striatal oscillations can emerge from inhibitory interactions between striatal medium spiny neurons (MSNs) (McCarthy *et al.*, 2011). This study suggests that the interaction of the synaptic gamma-aminobutyric acid currents and the intrinsic membrane M-current amplify oscillatory activities which can induce 8-30 Hz in the network after dopamine loss. Also, increased levels of cholinergic drive have been suggested as a condition relevant to the parkinsonian striatum, lead to synchronized beta oscillations in the striatal model (Kondabolu *et al.*, 2016). This study reported that striatal cholinergic interneuron can mediate the emergence of exaggerated beta oscillations in the basal ganglia circuit and that can induce parkinsonian-like motor deficits in animal models. In addition, computational neural network model of MSNs and fast-spiking interneurons (FSIs) found that besides imbalance firing rate, strong beta-band oscillations can emerge from implementation of changes to cellular and circuit properties caused by dopamine loss (Damodaran *et al.*, 2015). Drug-treatment to desynchronizing FSIs activities can reduce beta-band oscillations and imbalance in firing rate in the dopamine-depleted striatum (Damodaran *et al.*, 2015). A recent study suggests that in dopamine-depleted striatum, MSNs projecting to the indirect pathway, which exhibited increased firing rates and abnormal phase-locked firing to cortical beta oscillations, preferentially and excessively synchronized their firing at 15-30 Hz frequency band (Sharott *et al.*, 2017). These reports determine a cell-type-selective entrainment of striatal cells firing to parkinsonian beta oscillations. Altogether, these studies confirm the mechanistic role of striatal interneurons to induce synchronized beta (8-30 Hz) oscillations in the striatal network after dopamine depletion. In spite of above models and studies, there are still little-known facts about the role of specific cell types of the individual basal ganglia nuclei in the generation, propagation, and interaction of oscillatory dynamics throughout the basal ganglia circuits and its relation to motor function in PD.

Previous studies have addressed the current questions by comparing and contrasting the oscillations, but specifically in the 13-30 Hz in PD patients, MPTP-treated NHP models of PD, and 6-OHDA-lesioned rat (Sharott *et al.*, 2005; Hammond *et al.*, 2007; Stein and Bar-Gad, 2013). Similar to our study, a recent rodent study shows the evidence of the dopamine dependency of low-beta (14-20 Hz) directed interactions within the basal ganglia and cortico-basal ganglia networks (West *et al.*, 2018). The pharmacotherapy of PD is primarily based on two classes of drugs: dopamine precursors, (levodopa) and dopamine receptor agonists (apomorphine) and both therapies reduces the beta-band activity. However, levodopa is more effective in modulating beta-band power compared to other drugs (Kuhn *et al.*, 2017). In MPTP-treated NHP models, abnormal oscillations in the basal-ganglia network was more prominent in the lower frequency bands (<13 Hz) compared to PD patients (Devergnas *et al.*, 2014). This difference could be due to the disease model because dopamine cells die gradually over the time in human patients, however, in animal models, dopamine cells are died by local infusion of neurotoxin. Therefore, it has been proposed that cortico-basal ganglia circuits resonate with higher power in lower (<13 Hz) or beta frequency bands after dopamine loss in animal models. Here, we also found markedly increased striatal oscillations in the lower (8-13 Hz) and beta (13-20 Hz) frequency bands in MPTP-treated NHPs models compared to normal NHP. Since oscillatory power in the 8-20 Hz frequency band in the motor cortical and globus pallidal structures was similar to the striatal oscillatory power in the parkinsonian state, we suggest the presence of similar oscillatory dynamics within the cortico-basal ganglia circuitry. Consistent with previous results, the current study clearly demonstrates that dopamine regulates the generative mechanisms of 8-20 Hz oscillations in the striatum and downstream networks (Sharott *et al.*, 2005; Mallet *et al.*, 2008). Despite the well-defined association between 8-20 Hz oscillations and movement in the cortico-basal ganglia circuitry, the mechanisms underlying their generation remain elusive.

Increased 8-20 Hz oscillations have been confirmed in the extrastriatal basal ganglia nuclei and associated networks that appears after the dopamine loss (Singh, 2018). Based on our current results, we therefore confirm the occurrence of increased 8-20 Hz oscillatory activity in the striatum, main recipient of afferents to the basal ganglia, in parkinsonian models and the acute treatment of levodopa can normalize amplified 8-13 Hz and 13-20 Hz striatal oscillations. Consequently, for subsequent study, abnormal striatal oscillations should be interfered by means of different pharmacological interventions or electrical stimulation to affect the striatal mechanisms and downstream networks that underlie the bradykinetic features after dopamine loss in PD.

## Acknowledgements

This work was supported by NIH grants NS045962 and NS073994.

## References

Brown P. Oscillatory nature of human basal ganglia activity: relationship to the pathophysiology of Parkinson’s disease. Mov Disord 2003; 18(4): 357–63.

Brown P. Abnormal oscillatory synchronisation in the motor system leads to impaired movement. Curr Opin Neurobiol 2007; 17(6): 656–64.

Brown P, Oliviero A, Mazzone P, Insola A, Tonali P, Di Lazzaro V. Dopamine dependency of oscillations between subthalamic nucleus and pallidum in Parkinson’s disease. J Neurosci 2001; 21(3): 1033–8.

Costa RM, Lin SC, Sotnikova TD, Cyr M, Gainetdinov RR, Caron MG, et al. Rapid alterations in corticostriatal ensemble coordination during acute dopamine-dependent motor dysfunction. Neuron 2006; 52(2): 359–69.

Damier P, Hirsch EC, Agid Y, Graybiel AM. The substantia nigra of the human brain. II. Patterns of loss of dopamine-containing neurons in Parkinson’s disease. Brain 1999; 122 (Pt 8): 1437–48.

Damodaran S, Cressman JR, Jedrzejewski-Szmek Z, Blackwell KT. Desynchronization of fastspiking interneurons reduces beta-band oscillations and imbalance in firing in the dopamine-depleted striatum. J Neurosci 2015; 35(3): 1149–59.

de Hemptinne C, Swann NC, Ostrem JL, Ryapolova-Webb ES, San Luciano M, Galifianakis NB, et al. Therapeutic deep brain stimulation reduces cortical phaseamplitude coupling in Parkinson’s disease. Nat Neurosci 2015; 18(5): 779–86.

DeLong MR. Activity of pallidal neurons during movement. J Neurophysiol 1971; 34(3): 414–27.

Devergnas A, Pittard D, Bliwise D, Wichmann T. Relationship between oscillatory activity in the cortico-basal ganglia network and parkinsonism in MPTP-treated monkeys. Neurobiol Dis 2014; 68: 156–66.

Hammond C, Bergman H, Brown P. Pathological synchronization in Parkinson’s disease: networks, models and treatments. Trends Neurosci 2007; 30(7): 357–64.

Hutchison WD, Dostrovsky JO, Walters JR, Courtemanche R, Boraud T, Goldberg J, et al. Neuronal oscillations in the basal ganglia and movement disorders: evidence from whole animal and human recordings. J Neurosci 2004; 24(42): 9240–3.

Kondabolu K, Roberts EA, Bucklin M, McCarthy MM, Kopell N, Han X. Striatal cholinergic interneurons generate beta and gamma oscillations in the corticostriatal circuit and produce motor deficits. Proc Natl Acad Sci U S A 2016; 113(22): E3159–68.

Kuhn AA, Kupsch A, Schneider GH, Brown P. Reduction in subthalamic 8-35 Hz oscillatory activity correlates with clinical improvement in Parkinson’s disease. Eur J Neurosci 2006; 23(7): 1956–60.

Kuhn J, Haumesser JK, Beck MH, Altschuler J, Kuhn AA, Nikulin VV, et al. Differential effects of levodopa and apomorphine on neuronal population oscillations in the cortico-basal ganglia loop circuit in vivo in experimental parkinsonism. Exp Neurol 2017; 298(Pt A): 122–33.

Liang L, DeLong MR, Papa SM. Inversion of dopamine responses in striatal medium spiny neurons and involuntary movements. J Neurosci 2008; 28(30): 7537–47.

Litvak V, Jha A, Eusebio A, Oostenveld R, Foltynie T, Limousin P, et al. Resting oscillatory cortico-subthalamic connectivity in patients with Parkinson’s disease. Brain 2011; 134(Pt 2): 359–74.

Mallet N, Pogosyan A, Sharott A, Csicsvari J, Bolam JP, Brown P, et al. Disrupted dopamine transmission and the emergence of exaggerated beta oscillations in subthalamic nucleus and cerebral cortex. J Neurosci 2008; 28(18): 4795–806.

McCarthy MM, Moore-Kochlacs C, Gu X, Boyden ES, Han X, Kopell N. Striatal origin of the pathologic beta oscillations in Parkinson’s disease. Proc Natl Acad Sci U S A 2011; 108(28): 11620–5.

Potts LF, Wu H, Singh A, Marcilla I, Luquin MR, Papa SM. Modeling Parkinson’s disease in monkeys for translational studies, a critical analysis. Exp Neurol 2014; 256: 133–43.

Sharott A, Magill PJ, Harnack D, Kupsch A, Meissner W, Brown P. Dopamine depletion increases the power and coherence of beta-oscillations in the cerebral cortex and subthalamic nucleus of the awake rat. Eur J Neurosci 2005; 21(5): 1413–22.

Sharott A, Vinciati F, Nakamura KC, Magill PJ. A Population of Indirect Pathway Striatal Projection Neurons Is Selectively Entrained to Parkinsonian Beta Oscillations. J Neurosci 2017; 37(41): 9977–98.

Shimamoto SA, Ryapolova-Webb ES, Ostrem JL, Galifianakis NB, Miller KJ, Starr PA. Subthalamic nucleus neurons are synchronized to primary motor cortex local field potentials in Parkinson’s disease. J Neurosci 2013; 33(17): 7220–33.

Singh A. Oscillatory activity in the cortico-basal ganglia-thalamic neural circuits in Parkinson’s disease. Eur J Neurosci 2018.

Singh A, Liang L, Kaneoke Y, Cao X, Papa SM. Dopamine regulates distinctively the activity patterns of striatal output neurons in advanced parkinsonian primates. J Neurophysiol 2015; 113(5): 1533–44.

Singh A, Mewes K, Gross RE, DeLong MR, Obeso JA, Papa SM. Human striatal recordings reveal abnormal discharge of projection neurons in Parkinson’s disease. Proc Natl Acad Sci U S A 2016; 113(34): 9629–34.

Stein E, Bar-Gad I. beta oscillations in the cortico-basal ganglia loop during parkinsonism. Exp Neurol 2013; 245: 52–9.

West TO, Berthouze L, Halliday DM, Litvak V, Sharott A, Magill PJ, et al. Propagation of Beta/Gamma Rhythms in the Cortico-Basal Ganglia Circuits of the Parkinsonian Rat. J Neurophysiol 2018.

